# Systematic Description of 3q29 Duplication Syndrome Reveals New Syndromic Phenotypes: Results from the 3q29 Registry

**DOI:** 10.1101/697714

**Authors:** Rebecca M Pollak, Michael C Zinsmeister, Melissa M Murphy, Michael E Zwick, the Emory 3q29 Project, Jennifer G Mulle

## Abstract

3q29 duplication syndrome (3q29Dup) is a rare genomic disorder caused by the reciprocal duplication of the 1.6 Mb 3q29 deletion syndrome region. Case reports indicate the 3q29Dup is likely to be pathogenic, but because no systematic study of the syndrome exists, the full range of manifestations is not well-understood. To develop a better understanding of 3q29 duplication syndrome, we used the 3q29 registry (https://3q29deletion.patientcrossroads.org/) to ascertain 31 individuals with 3q29Dup, the largest cohort ever surveyed in a systematic way. For comparison, we ascertained 117 individuals with the reciprocal 3q29 deletion syndrome (3q29Del) and 64 typically developing controls. We used a custom medical and demographic questionnaire to assess physical and developmental phenotypes, and two standardized instruments, the Social Responsiveness Scale (SRS) and Achenbach Behavior Checklists (CBCL/ABCL), to assess social disability. We find that our 3q29Dup participants report a high rate of problems in the first year of life (80.6%), including feeding problems (58%), failure to gain weight (42%), hypotonia (39%), and respiratory distress (29%). In early childhood, learning problems (87.1%) and seizures (25.8%) are common. Additionally, we find a rate of self-reported ASD diagnoses (39%) similar to that previously identified in 3q29Del (29%), and the granular characteristics of social disability measured using the SRS and CBCL/ABCL are comparable between 3q29Dup and 3q29Del. This is the most comprehensive description of 3q29Dup to date. Our findings can be used to develop evidence-based strategies for early intervention and management of 3q29 duplication syndrome.

## INTRODUCTION

3q29 duplication syndrome (3q29Dup) is a genomic disorder caused by the reciprocal duplication of the 1.6 Mb 3q29 deletion syndrome (3q29Del) region (GRCh38 chr3: 195,998,129 – 197,623,129). This interval contains 21 distinct protein-coding genes, 3 antisense transcripts, 3 long noncoding RNAs, and 1 microRNA (Ballif et al., 2008; Willatt et al., 2005). 3q29Dup has been observed as both an inherited and *de novo* event (Ballif et al., 2008; Goobie et al., 2009; Vitale et al., 2018). The estimated prevalence of 3q29Dup from population-based studies ranges from ∼1:100,000 to ∼1:30,000 (Kaminsky et al., 2011; Männik et al., 2015; Moreno-De-Luca et al., 2013; Owen et al., 2018; Stefansson et al., 2014). Studies of individuals from clinical cohorts referred for microarray testing indicate a prevalence of ∼1:2000, a clear enrichment of the 3q29 duplication above population-based estimates (Ballif et al., 2008; Dittwald et al., 2013; Kaminsky et al., 2011; Moreno-De-Luca et al., 2013). However, the phenotype of 3q29Dup is not fully understood. Case reports have found 3q29Dup to be associated with developmental delay, speech delay, ID, ocular and cardiac anomalies, microcephaly, dental anomalies, obesity, and seizures (Aleixandre Blanquer et al., 2011; Ballif et al., 2008; Fernández-Jaén et al., 2014; Goobie et al., 2009; Kessi et al., 2018; Lesca et al., 2012; Lisi et al., 2008; Schilter et al., 2013; Tassano et al., 2018; Vitale et al., 2018). Additionally, some case reports have described behavioral phenotypes associated with 3q29Dup, most notably a disruptive behavioral profile (Quintela et al., 2015) and behavioral similarities to ASD (Lesca et al., 2012), and one case report identifies an individual with spina bifida (Lawrence, Arreola, Cools, Elton, & Wood, 2017). The largest case series published to date is a report on 19 individuals with 3q29Dup by Ballif et al (Ballif et al., 2008). Only five of these individuals had the canonical 1.6 Mb duplication; the other 14 cases had duplications of sizes varying from 200 kb to 2.4 Mb. Of the 19 total cases, seven had clinical information (Ballif et al., 2008). For the three subjects with the canonical 1.6 Mb duplication and clinical information, the only common feature was mild/moderate intellectual disability (Ballif et al., 2008).

Because the 3q29 duplication can be inherited from apparently unaffected parents, and because there is little data in the literature, there is no clear conclusion about the clinical significance of the duplication. For example, in ClinVar (https://www.ncbi.nlm.nih.gov/clinvar), of 19 submission entries for the 1.6 Mb 3q29 duplication, the variant is classified as pathogenic 13 times (68%) with the remaining entries classified as “Uncertain significance” (n = 5) or with “conflicting data from submitters” (n = 1). This means that genetic testing labs looking at the identical variant may classify it differently, which is confusing for families and clinicians alike. To bridge this knowledge gap, we have created an internet-based registry for individuals with 3q29Dup and 3q29Del (https://3q29deletion.patientcrossroads.org/) and implemented standardized instruments for systematic ascertainment of self-reported phenotypes. Here we present results from 31 individuals, the largest cohort of individuals with 3q29Dup ever described. We also compare phenotypes to 117 individuals with 3q29Del, to identify shared or divergent phenotypes that may be present across the syndromes.

Developing a clearer understanding of the phenotypic spectrum of 3q29Dup is crucial for clinicians, caregivers, and the probands themselves, so that evidence-based interventions can be synthesized. Furthermore, these data may provide insight into the potential molecular mechanism and gene dosage effects that have a role in the 3q29 CNVs. Additionally, it will be of clinical utility to determine whether the high rates of neuropsychiatric diagnoses and social disability in 3q29Del (Glassford et al., 2016; Pollak et al., 2019) are shared or distinct from 3q29Dup phenotypes, as this will provide guidance for developing clinical standards of care for this understudied population.

## METHODS

### Sample

Individuals with 3q29Dup were ascertained through the internet-based 3q29 registry (https://3q29deletion.patientcrossroads.org/) as previously reported (Glassford et al., 2016). Briefly, at launch in 2013, information about the registry was emailed to health care providers, medical geneticists, genetic counselors, and support organizations; the registry is currently advertised via Google AdWords, where specific keywords were chosen to target the registry website in internet searches. Participant recruitment, informed consent and assent, and data collection are all performed through the registry website. In April 2019, a data freeze was implemented and existing records were securely downloaded and de-identified for analysis. After data cleaning (removing spam accounts, duplicate records, and individuals with additional significant genetic diagnoses), 31 3q29Dup registrants (48.4% male) were included in the present study, ranging in age from 0.3-52.2 years (mean = 10.0±10.8 years). 117 individuals with 3q29Del (55.6% male) were also obtained through the 3q29 registry, ranging in age from 0.1-41.0 years (mean = 9.4±8.0 years). Clinical diagnosis of 3q29Dup or 3q29Del was confirmed via review of clinical genetics reports and/or medical records. Data from typically developing group controls (n = 64, 51.6% male) ranging in age from 1.0-41.0 years (mean = 9.9±7.2 years) were obtained as a comparison group (Pollak et al., 2019). Description of the study sample can be found in Table 1. This study was approved by Emory University’s Institutional Review Board (IRB00064133).

**Table 1:** Characteristics of study participants with 3q29Dup and controls.

|  | 3q29Dup | 3q29Del | Control |
| --- | --- | --- | --- |
| Age, years (mean ± SD) | 10.0 ± 10.8 | 9.4 ± 8.0 | 10.5 ± 7.2 |
| Sex (n, %) |  |  |  |
| Male | 15 (48.4%) | 65 (55.6%) | 33 (51.6%) |
| Female | 16 (51.6%) | 52 (44.4%) | 31 (48.4%) |
| Race (n, %) |  |  |  |
| White | 27 (87.1%) | 103 (88.0%) | 41 (64.1%) |
| Black/African American | 0 (0%) | 1 (0.9%) | 13 (20.3%) |
| Other | 4 (12.9%) | 11 (9.4%) | 8 (12.5%) |
| Blank | 0 (0%) | 2 (1.7%) | 2 (3.1%) |

Demographic data collected from the custom Medical & Demographic Questionnaire completed by participants upon enrollment in the online 3q29 Registry.

### Questionnaires

Upon registration, the participant or his/her parent or caregiver completed a custom medical and demographic questionnaire. This questionnaire includes questions on the sex, birthdate, race, and ethnicity of the participant, as well as a detailed medical history covering seven domains of physical and mental development: birth history, development, ear/nose/throat, gastrointestinal, renal, oral/dental, and seizures/psychiatric (Glassford et al., 2016).

In additional to medical phenotypes, two standardized questionnaires were used to assess ASD-related symptomology and general behavioral problems in the participants. The Social Responsiveness Scale (SRS; preschool, school-age, and adult forms; n = 15 3q29Dup, 67 3q29Del, 56 controls) is a 65-item, 4 point Likert-scaled questionnaire designed to assess ASD-related symptoms along a normative continuum (Constantino & Todd, 2012). The Child Behavior Checklist (CBCL) and Adult Behavior Checklist (ABCL) are 100-, 113-, or 126-item (CBCL preschool, CBCL school-age, and ABCL, respectively; n = 15 3q29Dup, 65 3q29Del, 57 controls), 3 point Likert-scaled questionnaires designed to assess behavioral or developmental problems (Achenbach & Rescorla, 2001, 2003). Data from the CBCL and ABCL were pooled for analysis. All standardized questionnaires were adapted for the online 3q29 registry and were completed by the participant or parent/guardian of the participant upon registration. Some participants were not eligible to complete the standardized questionnaires because the proband was too young. Demographic characteristics of the respondents for each questionnaire can be found in Table S1, demonstrating that the average age and sex distribution of participants who completed the medical and demographic questionnaire was not different from the average age and sex distribution of participants who completed each standardized form.

### Analysis

Diagnoses and health problems from the medical history questionnaire were recoded for analysis as yes/no binary variables; global developmental delay/intellectual disability (GDD/ID) diagnosis was recoded as yes (reported diagnosis of global developmental delay and/or intellectual disability)/no. Birthweight is coded in 1 lb increments in the online 3q29 registry (https://3q29deletion.patientcrossroads.org/); the midpoint of the interval was assumed as the birth weight of the participant for analysis. Developmental milestones are coded in “bins” of time in the registry, it was assumed that the milestone was reached at the midpoint of the selected interval for analysis. For milestones marked as “more than 10 years”, it was assumed the participant achieved the milestone at the midpoint between 10 years and their age at registration. For participants who had not yet reached a developmental milestone, their data were treated as censored observations, where time in the study is recorded consistent with age at the time of entry into the registry. To compare birthweight between 3q29Dup cases, 3q29Del cases, and controls, linear regression and goodness-of-fit analysis were implemented using the stats R package (Team, 2008), controlling for sex, gestational age, and race. To compare reported diagnoses between 3q29Dup cases and 3q29Del cases, Fisher’s exact test and chi-squared tests were implemented using the stats R package (Team, 2008). To compare length of time spent in the hospital between 3q29Dup cases and controls, two sample t-test was implemented using the stats R package (Team, 2008). To compare rates of self-reported seizures and psychiatric diagnoses in 3q29Dup cases to population prevalence values, one-sample proportion tests with Yates’ continuity correction were implemented using the stats R package (Team, 2008). Data from standardized questionnaires were imported into R (Team, 2008) and were recoded and scored according to the publisher’s guidelines. To compare standardized questionnaire scores between 3q29Dup cases, 3q29Del cases, and controls, linear regression was implemented using the stats R package (Team, 2008), controlling for age, race, and sex. To compare scores in 3q29Dup participants to mean values reported for children with idiopathic ASD (Torske, Naerland, Oie, Stenberg, & Andreassen, 2017), one sample t-test was implemented using the stats R package (Team, 2008). Kaplan-Meyer time-to-event analysis for developmental milestones was implemented using the survival R package (Therneau, 2015). Figures were generated using the plotly, ggplot2, and VennDiagram R packages (Chen, 2018; Sievert et al., 2017; Wickham, 2009).

## RESULTS

### Birth weight

The average birth weight for 31 3q29Dup participants is 6.50 lbs, with an average gestational age of 38.1 weeks, as compared to an average birth weight of 7.6 lbs and average gestational age of 39.2 weeks in our 64 typically developing controls (Figure 1). After adjusting for gestational age, sex, and race, 3q29Dup participants weigh significantly less than controls (p = 0.005), with an effect size of −0.74, indicating that 3q29Dup participants, on average, weigh 0.74 lbs (11.84 oz) less than control participants at birth. Additionally, goodness-of-fit analysis shows that including genotype (3q29Dup vs. control) in the model fits the data significantly better than only including gestational age, sex, and race (p = 0.005, Table S2). Due to the fact that the racial makeup is not matched between 3q29Dup participants and controls, we restricted the analysis to those participants that self-identify as white. We find that the magnitude of the effect size for the 3q29 duplication increases slightly, to −0.83 (p = 0.001), indicating that within self-identified white registrants, babies with 3q29Dup weigh 0.83 lbs (13.28 oz) less at birth than typically developing controls.

**Figure 1.**
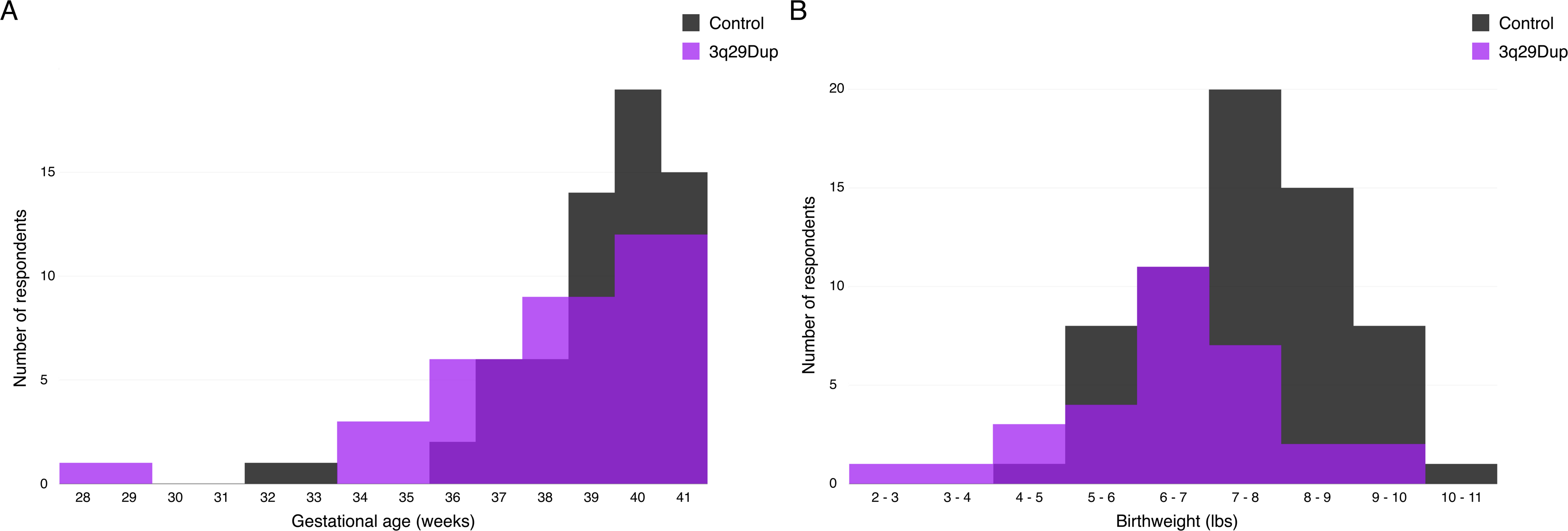
Gestational age and birthweight distributions for 3q29Dup and controls. **A)** Gestational age distribution for 3q29Dup participants (n = 31) and typically developing controls (n = 64). **B)** Birthweight distribution for 3q29Dup participants (n = 31) and typically developing controls (n = 64).

**Table 2:** SRS sub-score comparison stratified by genotype, sex, and ASD status.

|  | Social Awareness |  | Social Cognition |  | Social Communication |  | Social Motivation |  | RRB |  | SCI |  |
| --- | --- | --- | --- | --- | --- | --- | --- | --- | --- | --- | --- | --- |
|  | Mean ± SD | P value | Mean ± SD | P value | Mean ± SD | P value | Mean ± SD | P value | Mean ± SD | P value | Mean ± SD | P value |
| Genotype |  |  |  |  |  |  |  |  |  |  |  |  |
| Control | 47.04 ± 8.88 | - | 45.27 ± 7.64 | - | 45.88 ± 8.14 | - | 46.13 ± 7.66 | - | 47.66 ± 8.51 | - | 45.50 ± 7.74 | - |
| 3q29Dup | 72.53 ± 16.69 | 1.43E-10 | 74.53 ± 18.45 | 4.27E-13 | 77.27 ± 17.65 | 3.37E-14 | 71.13 ± 14.89 | 3.61E-12 | 81.47 ± 20.28 | 1.48E-13 | 77.27 ± 18.43 | 3.24E-14 |
| Sex |  |  |  |  |  |  |  |  |  |  |  |  |
| Male control | 45.97 ± 9.15 | - | 45.23 ± 7.42 | - | 45.73 ± 6.34 | - | 46.57 ± 6.58 | - | 47.63 ± 5.59 | - | 45.37 ± 6.60 | - |
| Male 3q29Dup | 71.83 ± 23.66 | 0.0003 | 78.67 ± 22.41 | 1.04E-06 | 78.83 ± 21.64 | 3.27E-07 | 73.33 ± 13.35 | 3.72E-07 | 85.83 ± 23.81 | 6.41E-08 | 79.67 ± 22.13 | 3.38E-07 |
| Female control | 48.27 ± 8.56 | - | 45.31 ± 8.03 | - | 46.04 ± 9.95 | - | 45.62 ± 8.85 | - | 47.69 ± 11.08 | - | 45.65 ± 9.01 | - |
| Female 3q29Dup | 73.00 ± 11.72 | 1.47E-07 | 71.78 ± 16.15 | 6.24E-07 | 76.22 ± 15.80 | 1.82E-07 | 69.67 ± 16.45 | 6.74E-06 | 78.56 ± 18.47 | 9.13E-07 | 75.67 ± 16.76 | 1.71E-07 |
| ASD status |  |  |  |  |  |  |  |  |  |  |  |  |
| Control | 47.04 ± 8.88 | - | 45.27 ± 7.64 | - | 45.88 ± 8.14 | - | 46.13 ± 7.66 | - | 47.66 ± 8.51 | - | 45.50 ± 7.74 | - |
| No ASD diagnosis 3q29Dup | 68.22 ± 17.09 | 1.74E-06 | 70.33 ± 19.66 | 1.50E-08 | 72.00 ± 18.73 | 3.56E-09 | 67.11 ± 16.83 | 1.00E-07 | 75.56 ± 21.37 | 1.29E-08 | 72.00 ± 19.72 | 3.31E-09 |
| ASD diagnosis 3q29Dup | 79.00 ± 15.15 | 5.86E-09 | 80.83 ± 15.99 | 1.24E-10 | 85.17 ± 13.64 | 2.85E-12 | 77.17 ± 9.77 | 3.80E-10 | 90.33 ± 16.23 | 7.50E-12 | 85.17 ± 14.27 | 3.15E-12 |

Comparison of mean scores on the SRS sub-scales between study participants with 3q29Dup and controls. 3q29Dup participants were stratified by sex and ASD status for further analysis. P values were calculated using simple linear regression, adjusting for age, race, and sex.

### Problems in the first year of life

3q29Dup participants spent longer in the hospital immediately after birth, with an average stay of 9.8 days (±14.5 days) as compared to an average of 3.8 days (±6.4 days) for typically developing controls (p = 0.037). Consistent with this longer hospital stay, 80.6% of 3q29Dup participants (n = 25) reported significant health problems in the first year of life, as compared to 39.1% of controls (n = 25). Some of these problems include: feeding problems (58.4%, n = 17); failure to gain weight (41.9%, n = 13); hypotonia (38.7%, n = 12); and respiratory distress (29.0%, n = 9). More data on problems reported in the first year of life can be found in Figure 2 and Table S3.

**Figure 2.**
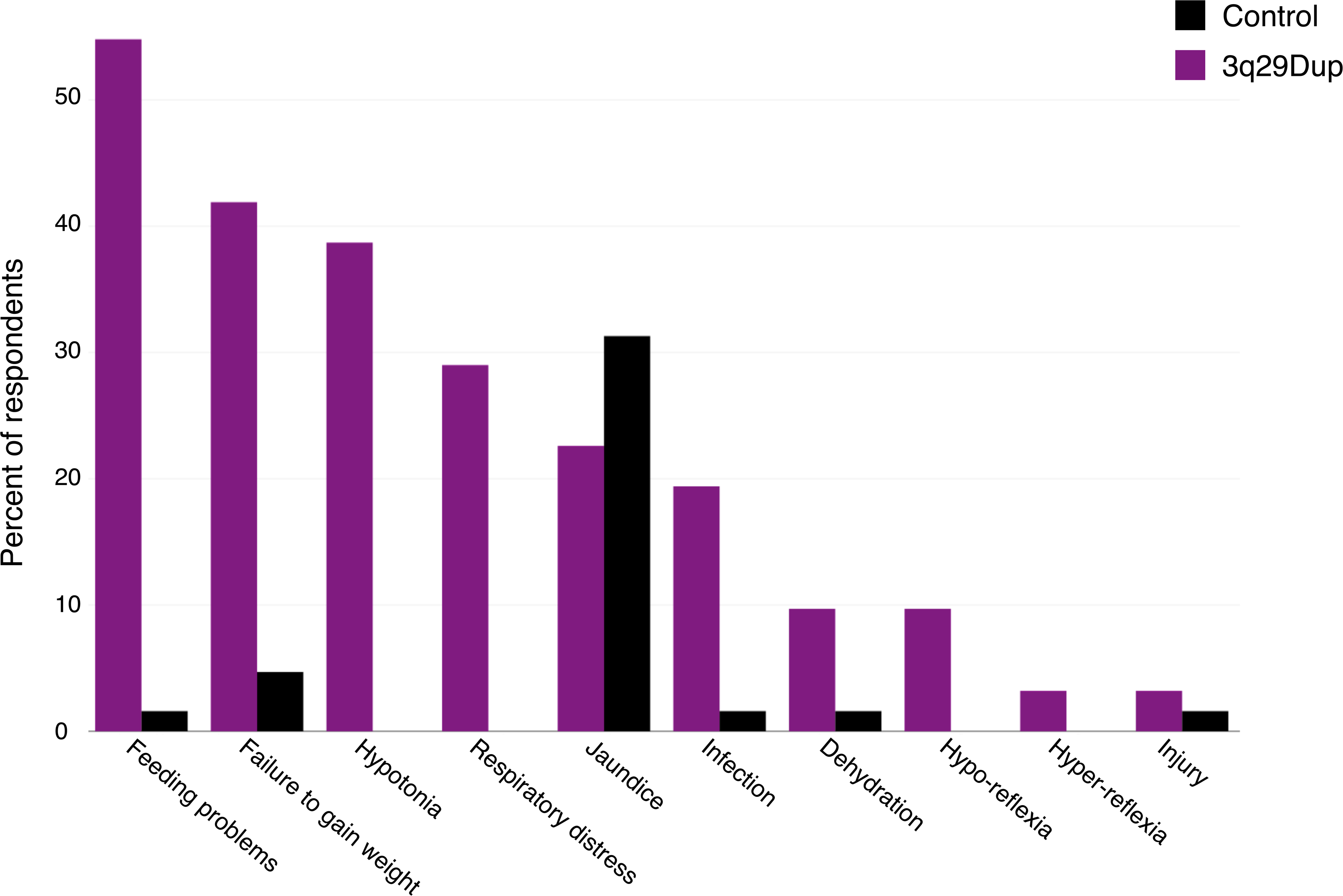
Reported problems in the first year of life by 3q29Dup participants and controls. Rate of problems in the first year of life reported by 3q29Dup participants (n = 31) and typically developing controls (n = 64), showing that 3q29Dup participants report substantially more problems in the postnatal period.

**Table 3:** CBCL/ABCL Withdrawn sub-score comparison stratified by genotype, sex, and ASD status.

| | Mean ± SD | P value |
| --- | --- | --- |
| Genotype |  |  |
| Control | 52.37 ± 5.85 | - |
| 3q29Dup | 66.20 ± 12.76 | 6.33E-08 |
| Sex |  |  |
| Male control | 52.14 ± 3.89 | - |
| Male 3q29Dup | 65.00 ± 9.96 | 5.56E-05 |
| Female control | 52.59 ± 7.33 | - |
| Female 3q29Dup | 67.00 ± 14.87 | 0.0004 |
| ASD status |  |  |
| Control | 52.37 ± 5.85 | - |
| No ASD diagnosis 3q29Dup | 64.88 ± 13.91 | 4.42E-05 |
| ASD diagnosis 3q29Dup | 67.71 ± 12.20 | 6.06E-06 |

Comparison of mean scores on the CBCL/ABCL Withdrawn sub-scale between study
participants with 3q29Dup and controls. 3q29Dup participants were stratified by sex and ASD status for further analysis. P values were calculated using simple linear regression, adjusting for age, race, and sex.

### Delay of developmental milestones

In the 3q29 registry, data are collected on social-emotional, communication, gross motor, and fine motor developmental milestones (Glassford et al., 2016). We used survival analysis to estimate the average time-to-event for developmental milestones for 3q29Dup participants and typically developing controls. One representative milestone was selected for each category; time-to-event curves for 3q29Dup and control participants are shown in Figure 3. For each milestone shown, 3q29Dup participants achieved that milestone on average 10 to 25 months later than typically developing controls (p <0.005); however, the majority of participants do eventually achieve each milestone. A full account of all milestones investigated is available in Table S4. Interestingly, while social-emotional, gross motor, and fine motor milestones on average are delayed by a similar amount (10 months, 15 months, and 16 months, respectively), communication milestones are more substantially delayed in 3q29Dup participants, with an average delay of 25 months.

**Figure 3.**
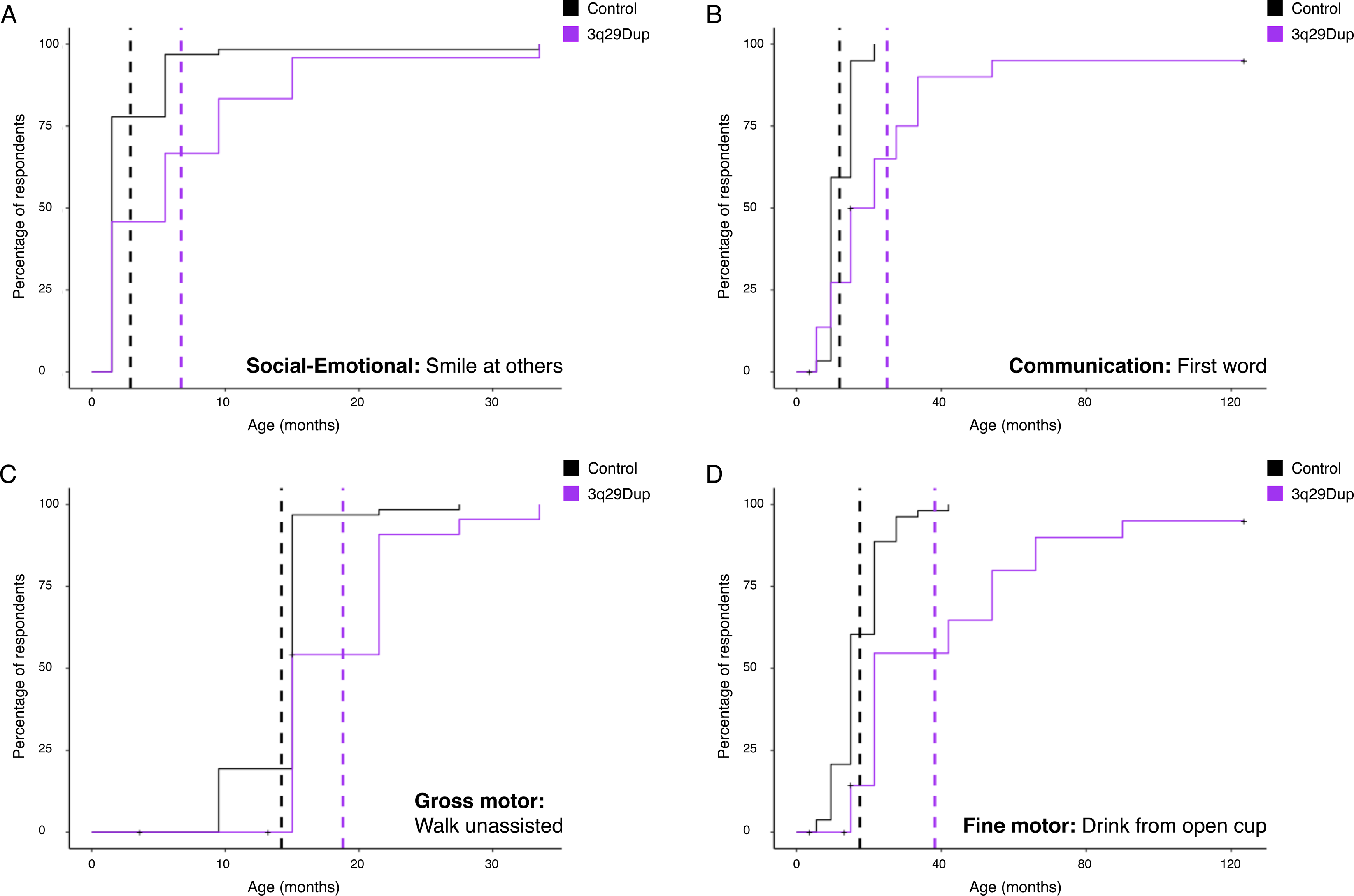
Comparison of developmental milestone achievement between 3q29Dup participants and controls. **A)** Kaplan-Meier time-to-event analysis of the representative social-emotional milestone, smile at others, showing that 3q29Dup participants (n = 24) on average achieve this milestone 3.79 months later than typically developing controls (n = 63). **B)** Kaplan-Meier time-to-event analysis of the representative communication milestone, first verbal word, showing that 3q29Dup participants (n = 23) on average achieve this milestone 13.4 months later than typically developing controls (n = 59). **C)** Kaplan-Meier time-to-event analysis of the representative gross motor milestone, walk unassisted, showing that 3q29Dup participants (n = 26 on average achieve this milestone 4.56 months later than typically developing controls (n = 62). **D)** Kaplan-Meier time-to-event analysis of the representative fine motor milestone, hold and drink from open cup, showing that 3q29Dup participants (n = 23) on average achieve this milestone 23.29 months later than typically developing controls (n = 53).

**Table 4:** Comparison of reported medical and neuropsychiatric phenotypes between 3q29Dup and 3q29Del participants.

| Category | 3q29Dup (% , n) | 3q29Del (% , n) | P value |
| --- | --- | --- | --- |
| Problems in the first year of life | 80.6% (25) | 82.1% (96) | 1.00 |
| Heart defects | 6.5% (2) | 24.8% (29) | 0.047 |
| Learning problems | 87.1% (27) | 79.5% (93) | 0.482 |
| GDD/ID | 45.2% (14) | 56.4% (66) | 0.360 |
| Ear problems | 41.9% (13) | 35.9% (42) | 0.682 |
| Gastrointestinal problems | 54.8% (17) | 67.5% (79) | 0.270 |
| Genitourinary problems | 6.5% (2) | 12.8% (15) | 0.527 |
| Renal problems | 16.1% (5) | 9.4% (11) | 0.329 |
| Dental problems | 67.7% (21) | 62.4% (73) | 0.734 |
| Seizures | 25.8% (8) | 15.4% (18) | 0.276 |
| Psychiatric diagnoses | 32.3% (10) | 35.0% (41) | 0.938 |

Comparison of medical and neuropsychiatric diagnosis categories collected from the custom
Medical & Demographic Questionnaire completed by participants upon enrollment in the online
3q29 Registry. P values were calculated using chi-squared test or Fisher's exact test.

### Learning disabilities

One of the most common phenotypes reported in prior studies of 3q29Dup is mild to moderate ID (Aleixandre Blanquer et al., 2011; Ballif et al., 2008; Lisi et al., 2008; Tassano et al., 2018), with one case report of a child with severe ID (Fernández-Jaén et al., 2014). Further, developmental delay, speech delay, and learning disabilities have been reported in individuals with 3q29Dup (Goobie et al., 2009; Quintela et al., 2015; Tassano et al., 2018), suggesting that neurodevelopmental and learning disabilities are common to individuals with 3q29Dup. Indeed, in our study population 87.1% (n = 27) of participants report at least one diagnosed learning problem, as compared to 4.7% (n = 3) of controls. Commonly reported learning problems include expressive language delay (54.8%, n = 17); global developmental delay (41.9%, n = 13); receptive language delay (29.0%, n = 9); learning disability in math (25.8%, n = 8); and learning disability in reading (25.8%, n = 8). A full account of learning disabilities reported by 3q29Dup and control participants can be found in Figure 4 and Table S5.

**Figure 4.**
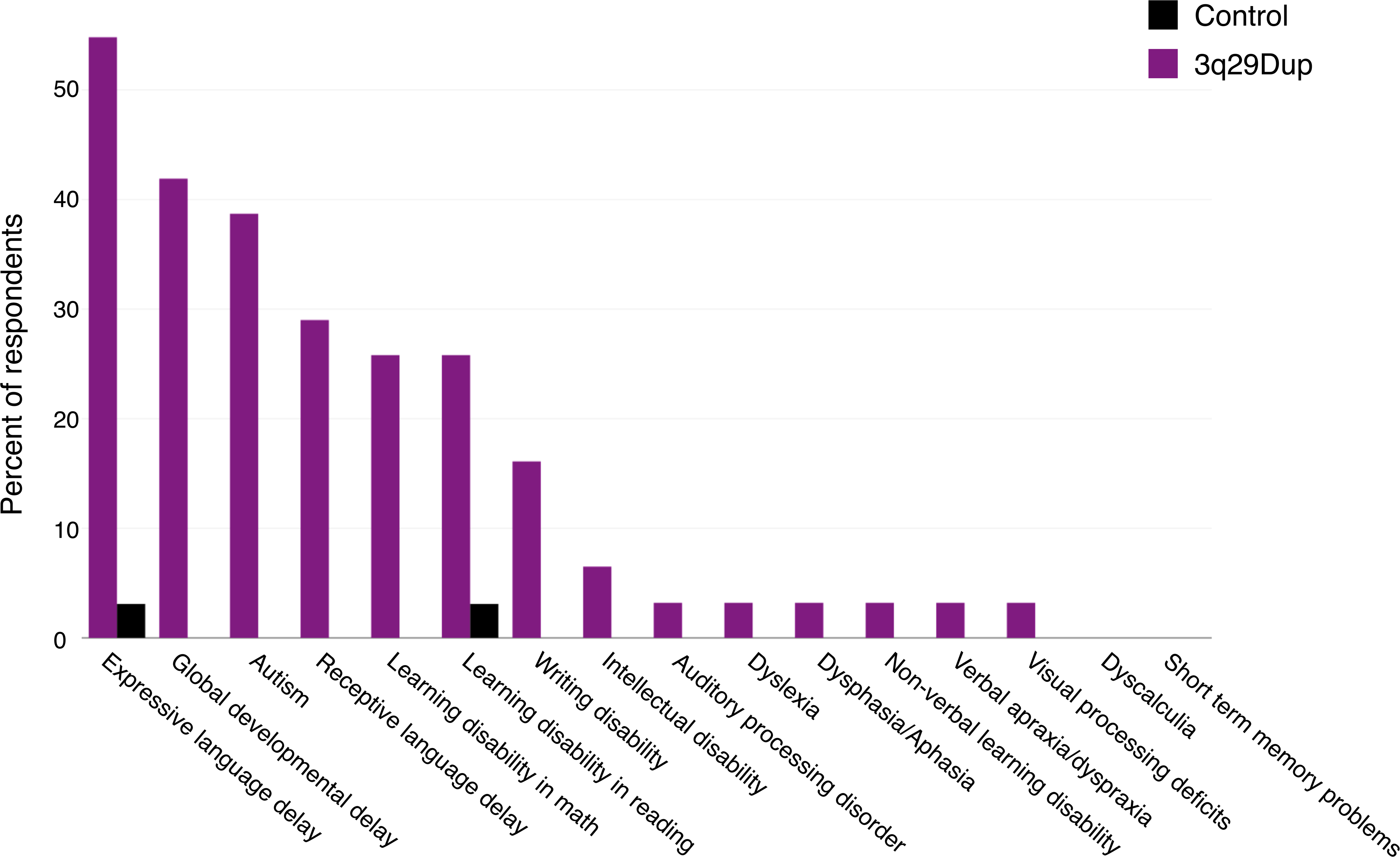
Reported learning problems by 3q29Dup participants and controls. Rate of learning problems reported by 3q29Dup participants (n = 31) and typically developing controls (n = 64), showing that 3q29Dup participants report substantially more learning problems.

### Gastrointestinal phenotypes

While gastrointestinal problems have not been previously reported in individuals with 3q29Dup, we find that 54.8% (n = 17) of 3q29Dup participants report at least one gastrointestinal problem (Table S6), including feeding problems beyond the first year of life (38.7%, n = 12) and chronic constipation (35.5%, n = 11).

### Seizures and neuropsychiatric phenotypes

*Seizures:* 25.8% (n = 8) of 3q29Dup participants reported seizures, consistent with prior reports of individuals with 3q29Dup (Fernández-Jaén et al., 2014; Kessi et al., 2018; Lesca et al., 2012; Tassano et al., 2018) and significantly elevated relative to the general population (general population prevalence = 1.2%, p < 2.20E-16) (Zack & Kobau, 2017).

*Neuropsychiatric diagnosis:* 32.3% (n = 10) of 3q29Dup participants reported a diagnosis of anxiety disorder (n = 10), significantly higher than the general population lifetime prevalence (5.7% vs. 32.3%, p = 1.05E-09). 38.7% (n = 12) of our 3q29Dup participants report a clinical diagnosis of ASD, a rate substantially higher than both that reported for 3q29Dup in the literature to date and that reported for the general population, with an estimated 26-fold increased risk for ASD for individuals with 3q29Dup. Subsets of 3q29Dup participants also report conduct disorder (6.5%, n = 2), depression (16.1%, n = 5), oppositional defiant disorder (6.5%, n = 2), and panic attacks (6.5%, n = 2) (Table S7). Additionally, there is only partial overlap between 3q29Dup participants reporting global developmental delay, ASD, seizures, and anxiety, indicating that these inflated rates are not due to a subset of severely affected participants, but rather are due to increased risks for these disorders in 3q29Dup (Figure 5).

**Figure 5.**
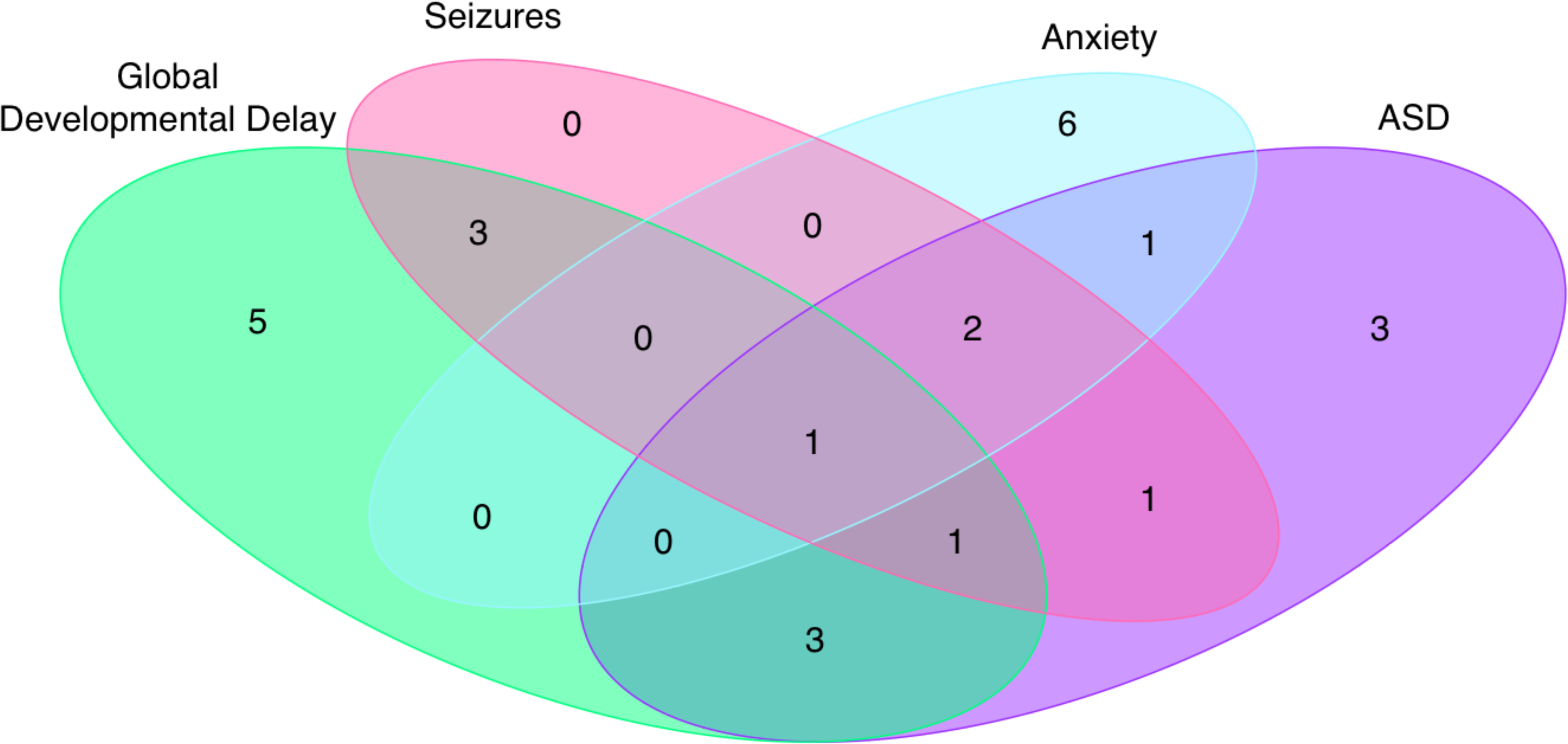
Overlap between reported global developmental delay, seizures, anxiety, and ASD among 3q29Dup participants. Venn diagram showing the overlap between reported global developmental delay, seizures, anxiety, and ASD within our 3q29Dup study population, demonstrating that these diagnoses are distributed through the population rather than clustered in a small group of severely affected participants.

### SRS social disability phenotypes and ASD features

While developmental delay and behavioral similarities to ASD have been identified in 3q29Dup (Goobie et al., 2009; Lesca et al., 2012; Tassano et al., 2018), social disability phenotypes have not been quantitatively described. Using standardized self-report tools, we find that participants with 3q29Dup have significantly higher total SRS scores than typically developing controls (3q29Dup mean *T-score* = 79.1, control mean *T-score* = 45.9, p = 1.09E-15). We observe this increased burden of social disability across sexes and ASD status within our 3q29Dup participants; individuals with 3q29Dup score significantly higher than controls irrespective of sex (3q29Dup female mean *T-score* = 77.1, control female mean *T-score* = 46.0, p = 2.22E-07; 3q29Dup male mean *T-score* = 82.0, control male mean *T-score* = 45.8, p = 1.83E-07) (Figure 6A) and ASD status (3q29Dup with ASD mean *T-score* = 87.2, 3q29Dup without ASD mean *T-score* = 73.7, control mean *T-score* = 45.9, p < 2.93E-09) (Figure 6B) (Table S8).

**Figure 6.**
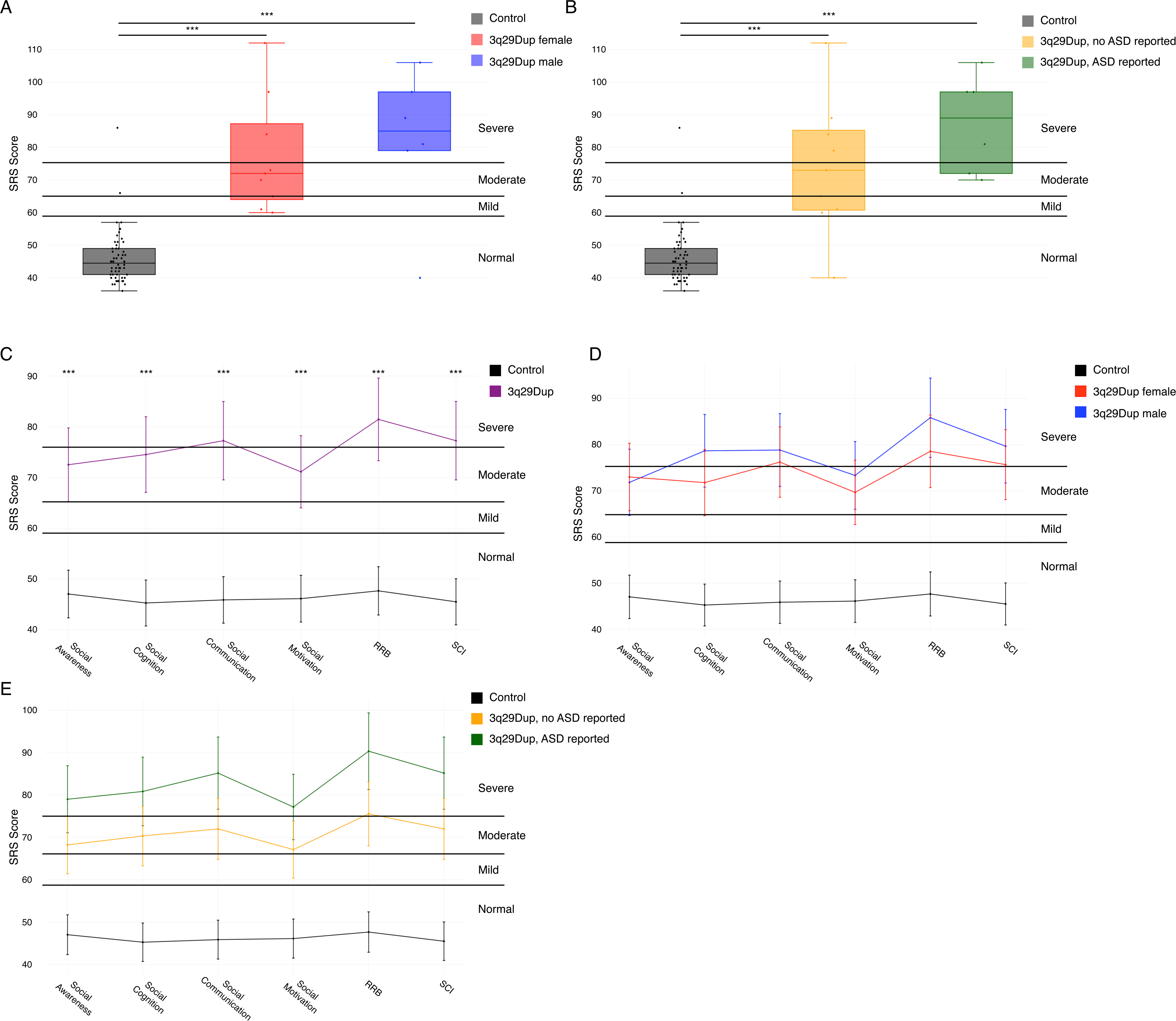
Comparison of SRS total scores and sub-scores between 3q29Dup and controls. **A)** SRS total scores split by controls (n = 56), 3q29Dup females (n = 9), and 3q29Dup males (n = 6), showing that 3q29Dup participants score significantly higher than controls irrespective of sex. **B)** SRS total scores split by controls (n = 56), 3q29Dup not reporting an ASD diagnosis (n = 9), and 3q29Dup reporting an ASD diagnosis (n = 6), showing that 3q29Dup participants score significantly higher than typically developing controls irrespective of self-reported ASD diagnosis status. **C)** Profile of individuals with 3q29Dup (n = 15) and controls (n = 56) across SRS sub-scales, showing moderate to severe impairment of 3q29Dup participants in all domains (RRB, Restricted Interests and Repetitive Behaviors; SCI, Social Communication and Interaction). **D)** Profile of 3q29Dup females (n = 9) and males (n = 6) and controls (n = 56) across SRS sub-scales, showing that 3q29Dup males and females both score significantly higher than controls and that the overall shape of the profile is consistent between 3q29Dup males and females. **E)** Profile of 3q29Dup participants reporting an ASD diagnosis (n = 6) and participants not reporting an ASD diagnosis (n = 9) and controls (n = 56) across SRS sub-scales, showing that 3q29Dup participants score significantly higher than controls irrespective of ASD status. ***, p < 0.001

#### SRS sub-scale profile in 3q29Dup

While the SRS total score can give an indication of the overall degree of social impairment for an individual, the SRS sub-scales can provide more detail about specific domains of social functioning that may be compromised. We analyzed all SRS sub-scales (Social Awareness, Social Cognition, Social Communication, Social Motivation, Restricted Interests and Repetitive Behaviors, and Social Communication and Interaction) to determine whether the inflated SRS total scores we observed are attributable to substantial deficits in all domains or functioning, or if 3q29Dup individuals show specific impairments in a few domains. Mean scores for Social Communication (*T-score* = 77.3), Restricted Interests and Repetitive Behaviors (*T-score* = 81.5), and Social Communication and Interaction (*T-score* = 77.3) were in the severe range, while mean scores for Social Awareness (*T-score* = 72.5), Social Cognition (*T-score* = 74.5), and Social Motivation (*T-score* = 71.1) were in the moderate range (Figure 6C, Table 2). Notably, 3q29Dup participants have significantly lower Social Motivation scores than those reported in cases of idiopathic ASD (3q29Dup Social Motivation *T-score* = 71.1, idiopathic ASD *T-score* = 78.4, p = 0.040) (Torske et al., 2017), indicating a preservation of social motivation relative to other domains assessed by the SRS. 3q29Dup participants scored significantly higher than typically developing controls on all sub-scales (p < 4.0E-11) (Table 2).

#### SRS sub-scale profile stratified by sex

We find that males and females with 3q29Dup do not score significantly differently from each other on any sub-scale (p > 0.5); however, both males and females with 3q29Dup score significantly higher than controls (p < 0.0005, Table 2), similar to our previous finding in 3q29Del (Pollak et al., 2019). Interestingly, 3q29Dup females score approximately 3 points higher than males on the Social Awareness sub-scale, while 3q29Dup males score slightly higher than females on all other sub-scales (Figure 6D), suggesting that, with larger sample size, some differences in social disability may emerge between males and females with 3q29Dup.

#### SRS sub-scale profile stratified by ASD diagnosis

Similar to 3q29Del (Pollak et al., 2019), we find that the shape of the SRS sub-score profile is shared between 3q29Dup participants with and without ASD, with 3q29Dup participants reporting an ASD diagnosis scoring on average 10 to 15 points higher on every sub-scale than 3q29Dup participants not reporting an ASD diagnosis (Figure 6E). Consistent with 3q29Dup participants having higher total SRS scores than controls irrespective of ASD status, 3q29Dup participants also score in the moderate or severe range, and significantly higher than controls, on all SRS sub-scales irrespective of ASD status (p < 2.0E-06, Table 2).

### CBCL/ABCL behavioral phenotypes

#### CBCL/ABCL Withdrawn sub-scale

To further investigate social disability phenotypes in 3q29Dup, we used the Withdrawn sub-scale of the CBCL and ABCL. Previous studies have shown that individuals with idiopathic ASD, on average, score in the borderline range on this sub-scale, and the majority of individuals score in the borderline or critical range (Mazefsky, Anderson, Conner, & Minshew, 2011; Noterdaeme, Minow, & Amorosa, 1999). Here, we find that 3q29Dup participants score significantly higher than controls (3q29Dup mean *T-score* = 66.2, control mean *T-score* = 52.4; p < 4.0E-07; Table 3). The average score for 3q29Dup participants overall, and males and females separately, is in the borderline range (Figure 7A and B, Table 3). This supports the SRS data and suggests a previously unidentified social disability phenotype in 3q29Dup. Over 50% of 3q29Dup participants score in the borderline or clinical range, with similar proportions observed in participants reporting a diagnosis of ASD and those not reporting a diagnosis of ASD (Figure 7C, Table 3).

**Figure 7.**
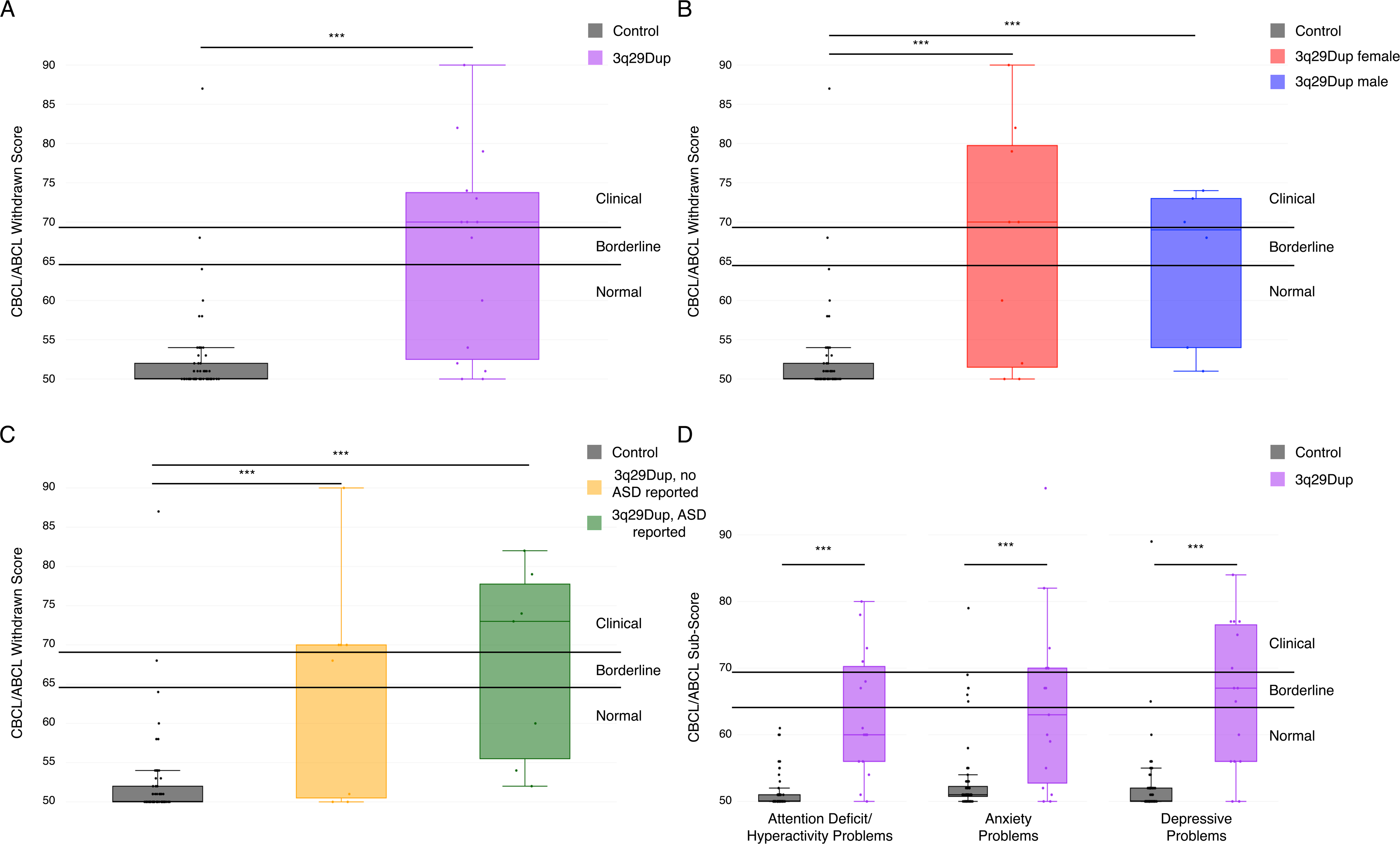
Comparison of CBCL/ABCL Withdrawn and DSM-oriented sub-scales between 3q29Dup and controls. **A)** Profile of 3q29Dup participants (n = 15) and controls (n = 57) on the Withdrawn sub-scale from the CBCL and ABCL, showing a significantly higher score in 3q29Dup participants, with a mean score in the moderate range. **B)** Profile of 3q29Dup males (n = 6) and females (n = 9) and controls (n = 57) on the Withdrawn sub-scale of the CBCL and ABCL, showing that both males and females score significantly higher than typically developing controls. **C)** Profile of 3q29Dup participants reporting an ASD diagnosis (n = 7) and not reporting an ASD diagnosis (n = 8) and controls (n = 57) on the Withdrawn sub-scale of the CBCL and ABCL, showing that 3q29Dup participants score significantly higher than typically developing controls irrespective of ASD status. **D)** Profile of 3q29Dup participants (n = 15) and controls (n = 57) across 3 DSM-oriented sub-scales from the CBCL and ABCL, showing significantly increased pathology in 3q29Dup participants across all domains. ***, p < 0.001

#### CBCL/ABCL DSM-oriented sub-scales

To assess additional behavioral features of 3q29Dup, we examined the DSM-oriented Attention Deficit/Hyperactivity Problems, Anxiety Problems, and Depressive Problems sub-scales from the CBCL and ABCL. 3q29Dup participants score significantly higher than controls on every sub-scale (3q29Dup Attention Deficit/Hyperactivity Problems *T-score* = 63.0, control Attention Deficit/Hyperactivity Problems *T-score* = 51.3; 3q29Dup Anxiety Problems *T-score* = 64.4, control Anxiety Problems *T-score* = 53.2; 3q29Dup Depressive Problems *T-score* = 65.8, control Depressive Problems *T-score* = 52.3; all p < 0.0005) (Figure 7D). These data suggest additional neuropsychiatric phenotypes are associated with the 3q29 duplication.

### Other phenotypes

Two participants (6%) reported heart defects (Table S9), two participants (6%) reported genitourinary phenotypes (Table S10), five participants (16%) reported renal phenotypes (Table S11), 13 (42%) participants reported ear problems (Table S12), 21 (68%) participants reported dental problems (Table S13), and one participant (3%) reported cleft palate (Table S14).

### Comparison of 3q29Dup and 3q29Del

*Medical phenotypes:* To determine whether there is evidence for divergent phenotypes associated with 3q29Dup and 3q29Del, we compared overall rates of reported problems in the first year of life, heart defects, learning problems, GDD/ID, ear problems, gastrointestinal problems, genitourinary problems, renal problems, dental problems, seizures, and psychiatric diagnoses between 3q29Dup and 3q29Del participants (Table 4). Congenital heart defects are reported at a significantly higher rate by 3q29Del participants as compared to 3q29Dup participants (24.8% vs. 6.5%, p = 0.047). Although not statistically significant, 3q29Dup participants reported seizures at a rate substantially greater than 3q29Del participants (25.8% vs. 15.4%, p = 0.276), and 3q29Del participants reported genitourinary problems at a rate twice that of 3q29Dup participants (12.8% vs. 6.5%, p = 0.527). Rates of all other reported problems were remarkably similar between 3q29Dup and 3q29Del participants, suggesting that the reciprocal 3q29 CNVs have similar effects on organ systems.

*Psychiatric phenotypes and social disability:* ASD is present at a similar frequency in both disorders (38.7% in 3q29Dup vs 29.1%% in 3q29Del, p = 0.416); however, anxiety disorder is slightly more commonly reported by 3q29Dup participants (32.3% in 3q29Dup vs 28.2% in 3q29Del, p = 0.048) (Table S15). The 3q29 deletion is established as an ASD-risk variant (Pollak et al., 2019; Sanders et al., 2015) but it is not known whether social disability phenotypes are similarly present in the 3q29 duplication. Using the SRS, we find that 3q29 Dup participants score similarly to 3q29Del participants (3q29Dup mean *T-score* = 79.1, 3q29Del mean *T-score* = 72.9, p = 0.107) (Figure 8A), indicating a similar burden of social disability shared between 3q29Dup and 3q29Del. The distribution of SRS sub-scores between 3q29 Dup and 3q29 Del is qualitatively similar (Figure 8B). However, 3q29Dup participants score significantly higher on Social Motivation than 3q29Del participants (3q29Dup Social Motivation *T-score* = 71.1, 3q29Del Social Motivation *T-score* = 63.5, p = 0.043), indicating that 3q29Dup cases have an intermediate social motivation phenotype, with significantly more impairment than that observed in 3q29Del, but significantly less impairment that that reported in idiopathic ASD (Torske et al., 2017). For the CBCL/ABCL Withdrawn sub-scale, we have previously reported that individuals with 3q29Del have mean scores in the normal range on this sub-scale, and that over 50% of 3q29Del individuals reporting a diagnosis of ASD score in the borderline or clinical range (Pollak et al., 2019). Here, we find that 3q29Dup and 3q29Del participants both score significantly higher than controls (3q29Dup mean *T-score* = 66.2, 3q29Del mean *T-score* = 63.2, control mean *T-score* = 52.4; p < 4.0E-07) and that they do not score significantly differently from each other (p = 0.318). However, the average score for 3q29Dup participants overall, and males and females separately, is in the borderline range (Figure 7A and B, Table 3), whereas the mean score for 3q29Del participants is in the normal range, suggesting a more substantial, and previously unidentified, social disability phenotype in 3q29Dup as compared to 3q29Del.

**Figure 8.**
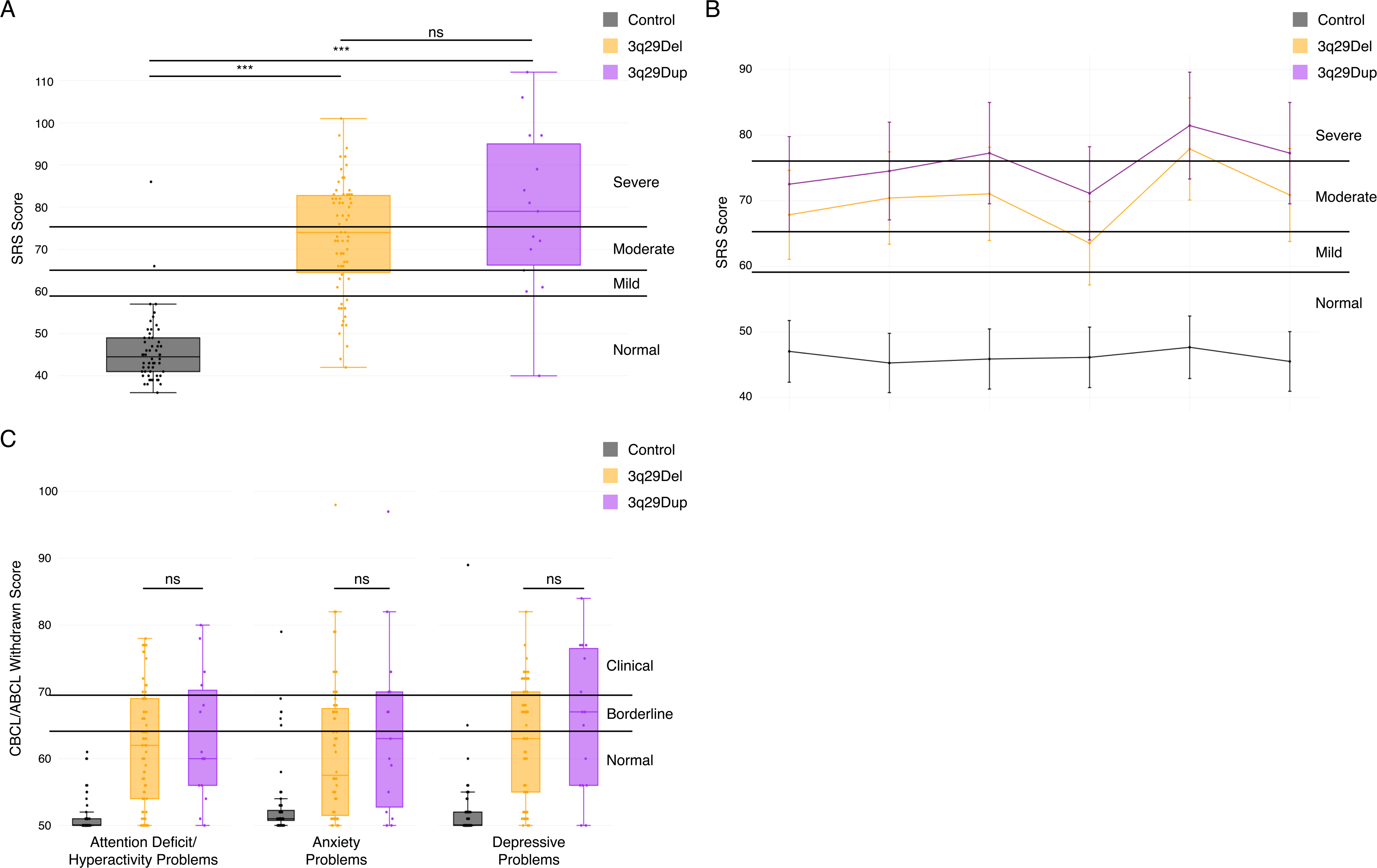
Comparison of SRS and CBCL/ABCL scores between 3q29Dup and 3q29Del. **A)** Total scores on the SRS for 3q29Dup participants (n = 15), 3q29Del participants (n = 67), and controls (n = 64), showing that 3q29Dup and 3q29Del participants do not score significantly differently. **B)** SRS sub-scale profile for 3q29Dup participants\ (n = 15), 3q29Del participants (n = 67), and controls (n = 64), showing that the shape of the SRS sub-scale profile is conserved between 3q29Dup and 3q29Del participants. **C)** Profile of 3q29Dup participants (n = 15), 3q29Del participants (n = 64), and controls (n = 57) across 3 DSM-oriented sub-scales from the CBCL and ABCL, showing that 3q29Dup and 3q29Del participants have similar levels of pathology in all 3 domains. ns, not significant.

To determine whether 3q29Dup shares some behavioral features with 3q29Del, we examined the DSM-oriented Attention Deficit/Hyperactivity Problems, Anxiety Problems, and Depressive Problems sub-scales from the CBCL and ABCL. Both 3q29Dup and 3q29Del participants score significantly higher than controls on every sub-scale (3q29Dup Attention Deficit/Hyperactivity Problems *T-score* = 63.0, 3q29Del Attention Deficit/Hyperactivity Problems *T-score* = 61.7, control Attention Deficit/Hyperactivity Problems *T-score* = 51.3; 3q29Dup Anxiety Problems *T-score* = 64.4, 3q29Del Anxiety Problems *T-score* = 61.4, control Anxiety Problems *T-score* = 53.2; 3q29Dup Depressive Problems *T-score* = 65.8, 3q29Del Depressive Problems *T-score* = 63.1, control Depressive Problems *T-score* = 52.3; all p < 0.0005), and 3q29Dup and 3q29Del participants do not score significantly differently from each other (p > 0.282, Figure 8C), suggesting that participants with 3q29Dup and 3q29Del have shared liability for these neuropsychiatric phenotypes previously associated with 3q29Del (Glassford et al., 2016).

## DISCUSSION

This study is the first to report on phenotypes associated with 3q29Dup using a systematic, standardized approach. We find a high prevalence of problems in the first year of life, including feeding problems, failure to gain weight, hypotonia, and respiratory distress, suggesting that individuals with 3q29 duplication require extra clinical attention during infancy. We also find that seizures, frequently described in case reports of 3q29 duplication syndrome (Fernández-Jaén et al., 2014; Kessi et al., 2018; Lesca et al., 2012; Tassano et al., 2018), are reported in 25% of our study subjects, thus individuals with 3q29 duplication syndrome should be evaluated by a pediatric neurologist. We find feeding problems and chronic constipation are reliably manifest, such that a pediatric gastroenterologist should administer an evaluation. Our data also suggest that ASD and social disability phenotypes are enriched in 3q29 duplication syndrome, and individuals with the 3q29 duplication should therefore be evaluated for ASD using gold-standard clinical measures.

We have found that the 3q29 duplication carriers report a substantial reduction in birthweight (0.74 lbs, 11.84 oz), similar to that previously reported for the reciprocal 3q29 deletion (Glassford et al., 2016). In 3q29 deletion syndrome, unpublished data from human subjects assessed by the Emory 3q29 Project (http://genome.emory.edu/3q29/) (Murphy et al., 2018) and existing data from the 3q29 mouse model (Baba et al., 2019; Rutkowski et al., 2019) show that the weight deficit in 3q29 deletion syndrome persists into adolescence. However, in an apparent paradox, the 3q29 duplication is associated with obesity (Ballif et al., 2008; Fernández-Jaén et al., 2014; Goobie et al., 2009; Lisi et al., 2008; Vitale et al., 2018), thus 3q29 duplication carriers weigh less at birth but may exhibit accelerated weight gain at an unknown developmental timepoint. Although the 3q29 registry does not collect data on current weight and height for study participants, these data suggest a compelling future direction for longitudinal data collection on weight and height. These findings also support a complex dose-response relationship between 3q29 interval genes and metabolic phenotypes.

Prior to this study, the 3q29 duplication had not been linked to ASD and related social disability phenotypes. One case study reported behavioral similarities to ASD (Lesca et al., 2012); however, 3q29Dup cases have not been identified in cohort studies of ASD (Moreno-De-Luca et al., 2013). It is possible that the population frequency of the 3q29 duplication is low, such that current studies of ASD are underpowered to find association with ASD. As larger cohorts become available, an association between ASD and the 3q29 duplication may become apparent. It is also possible that with existing genomics technologies (such as array-based methods), duplications are challenging to identify and are susceptible to high false-negative rates. Improved analysis methods or new technologies (Mohr et al., 2017) may rectify this problem. It is also possible that our study suffers from ascertainment bias, where individuals with ASD are referred for genetic testing and are coincidentally found to have a 3q29 duplication. However, our own data partially contradict this possibility. In figures 6 and 7, we show that substantial social disability is present even among individuals *without a diagnosis of ASD*, as assessed by both the SRS and the CBCL. Our data suggest the 3q29 duplication is a susceptibility locus for ASD, and individuals with the 3q29 duplication should be evaluated for ASD using gold-standard clinical measures.

We found that the rate of reported ASD diagnoses in our 3q29Dup study population was elevated as compared to 3q29Del, but the difference did not reach statistical significance (38.7% vs. 29.1%, p = 0.416). Because this is the first study to use standardized, quantitative measures to assess dimensions of social disability in 3q29Dup, we are able to evaluate nuances of social behavior in our study sample. Our findings suggest that the degree of social disability in this population has been underappreciated in the literature. Further, we found a similar profile of ASD features on the SRS to the profile we had previously identified in 3q29Del (Pollak et al., 2019), suggesting that the 3q29 CNVs harbor similar risk for social disability, irrespective of ASD diagnosis status. Previous work by our group found that 3q29Del individuals have relatively well-preserved social motivation as assess by the SRS Social Motivation sub-scale (Pollak et al., 2019); here, we report that 3q29Dup individuals appear to have an intermediate social motivation phenotype, with Social Motivation sub-scale scores significantly higher (more impaired) than 3q29Del, but significantly lower (less impaired) than those reported in a study of idiopathic ASD (Torske et al., 2017). Additionally, we found that 3q29Dup participants, on average, scored in the borderline range for the Withdrawn sub-scale of the CBCL and ABCL, which further supports a previously unappreciated degree of social disability in the 3q29Dup population. Taken together, this indicates that, similar to 3q29Del, individuals diagnosed with 3q29Dup should receive gold-standard ASD evaluations as a standard of care.

While we identified several similarities between 3q29Dup and 3q29Del, including the effect of the 3q29 CNV on birthweight, SRS scores, and CBCL/ABCL scores, there are also features where 3q29Dup and 3q29Del diverge. We found that there is a significantly higher rate of congenital heart defects in 3q29Del as compared to 3q29Dup (24.8% vs. 6.5%, p = 0.047), which is in line with previously published studies of 3q29Del and 3q29Dup. We also found a higher rate of 3q29Dup participants reporting seizures as compared to 3q29Del participants (25.8% vs. 15.4%, p = 0.276); while this difference is not statistically significant, it lends additional support to the association between 3q29Dup and seizure phenotypes previously described in case reports (Fernández-Jaén et al., 2014; Kessi et al., 2018; Lesca et al., 2012; Tassano et al., 2018). The registry asks only about the presence or absence of seizures; collecting more data about seizure phenotypes associated with the 3q29 duplication is an important future direction for this work. While the overall rate of individuals reporting at least one psychiatric diagnosis was similar between 3q29Dup and 3q29Del (32.3% vs. 35.0%, p = 0.938), the rate of each psychiatric disorder varies between 3q29Dup and 3q29Del (Table S15). As ongoing efforts to articulate the molecular mechanism and downstream targets impacted by the 3q29 duplication and 3q29 deletion bear fruit, it will be productive to compare convergent and divergent downstream pathways with the concordant and discordant phenotypic spectra of these reciprocal disorders.

Although there are significant strengths to this study, it is not without limitations. While this is the largest cohort of 3q29Dup individuals reported on to date, the small sample size restricts our ability to draw definitive conclusions due to decreased statistical power. We found substantial differences in some health domains, most notably seizures, that did not reach statistical significance due to our sample size. Studies with larger sample size will be better able to assess the importance of these differences between 3q29Dup and 3q29Del. Additionally, all of the data used in this study was collected using parent-report measures, which introduces the potential of recall bias due to the data being retrospective. However, a previous study by our group using the same measures as the present study found high concordance between parent-reported diagnoses and direct assessment (Pollak et al., 2019). Lastly, there are two potential sources of ascertainment bias in our study. First, our sample of 3q29Dup participants is overwhelmingly white, suggesting that we are not effectively reaching minority populations with our recruitment efforts. Second, we note that parents that register their children and complete time-consuming questionnaires are likely to be highly motivated, potentially because their children are severely affected. If our study sample is taken from the extreme end of 3q29Dup phenotypes, scores on the SRS and CBCL/ABCL and reported health problems and diagnoses are likely to be inflated as compared to the true prevalence in the 3q29Dup population. Direct assessment of individuals with 3q29Dup by the Emory 3q29 Project (http://genome.emory.edu/3q29/) (Murphy et al., 2018) aim to address some of the weakness of this work by performing comprehensive gold-standard evaluations by expert clinicians.

There are significant strengths of this study, most notably that we have reported on the largest cohort of individuals with 3q29Dup to date. Further, we have systematically ascertained phenotypes from our study population and from a population of individuals with 3q29Del, and we were able to compare phenotypes between the two groups. Although we had a relatively small sample size, we were able to identify significant differences between 3q29Dup and 3q29Del participants in the frequency of congenital heart defects, and we were able to find suggestive evidence of other phenotypic divergences between the reciprocal CNVs. We were also able to identify areas of phenotypic concordance between 3q29Dup and 3q29Del; if these similarities are borne out by studies with larger sample size, it could provide meaningful insight into the molecular mechanisms driving phenotype development, including neuropsychiatric and neurodevelopmental phenotypes. Finally, we found a substantial degree of social disability in our 3q29Dup population that has not been previously reported in the literature, which suggests that gold-standard ASD evaluations should be standard of care for individuals with 3q29Dup. Taken together, this study serves as a valuable complement to previously published case studies of 3q29Dup; by systematically ascertaining phenotypes and comparing them to 3q29Del and typically developing controls, we are able to add to the body of knowledge regarding the 3q29Dup phenotype and find relationships between 3q29Dup and 3q29Del, similar to those identified in other reciprocal CNV disorders. These data will assist clinicians, caregivers, and probands in developing comprehensive treatment plans to improve long-term outcomes for individuals with 3q29Dup.

## ACKNOWLEDGEMENTS

We gratefully acknowledge our study population, the 3q29 duplication community, for their participation and commitment to research.

We also acknowledge the contributions of the members of the Emory 3q29 Project: Hallie Averbach, Emily Black, Gary J Bassell, T Lindsey Burrell, Grace Carlock, Tamara Caspary, Joseph F Cubells, David Cutler, Paul A Dawson, Michael P Epstein, Roberto Espana, Michael J Gambello, Katrina Goines, Henry R Johnston, Cheryl Klaiman, Sookyong Koh, Elizabeth J Leslie, Longchuan Li, Bryan Mak, Tamika Malone, Trenell Mosley, Derek Novacek, Ryan Purcell, Timothy Rutkowski, Rossana Sanchez, Celine A Saulnier, Jason Schroeder, Esra Sefik, Brittney Sholar, Sarah Shultz, Nikisha Sisodiya, Steven Sloan, Elaine F Walker, Stephen T Warren, David Weinshenker, and Zhexing Wen.

## CONFLICTS OF INTEREST

The authors have no conflicts of interest to disclose.

