## Supplemental information for "Systematic Description of 3q29 Duplication Syndrome Reveals New Syndromic Phenotypes: Results from the 3q29 Registry"

**Table S1: Questionnaire demographics.** Characteristics of study participants with 3q29Dup and controls completing each questionnaire utilized in present study.

| | Medical and Demographic Questionnaire | | | Social Responsiveness Scale (SRS) | | | Achenbach Behavior Checklists (CBCL/ABCL) | | |
|---|---|---|---|---|---|---|---|---|---|
| | 3q29Dup | 3q29Del | Control | 3q29Dup | 3q29Del | Control | 3q29Dup | 3q29Del | Control |
| Age, years (mean ± SD) | 10.0 ± 10.8 | 9.4 ± 8.0 | 10.5 ± 7.2 | 11.5 ± 9.3 | 10.4 ± 7.3 | 10.9 ± 7.2 | 9.1 ± 6.2 | 9.7 ± 6.7 | 10.0 ± 7.0 |
| Sex (%, n) | | | | | | | | | |
| Male | 48.4% (15) | 55.6% (65) | 51.6% (33) | 40.0% (6) | 55.2% (37) | 53.6% (30) | 40% (6) | 56.3% (36) | 49.1% (28) |
| Female | 51.6% (16) | 44.4% (52) | 48.4% (31) | 60.0% (9) | 44.8% (30) | 46.4% (26) | 60.0% (9) | 43.8% (28) | 50.9% (29) |

**Table S2: Goodness-of fit analysis.** Goodness-of-fit analysis for testing the effect of genotype on birthweight, showing that a model including genotype fits the data significantly better than one without. Y is birthweight in pounds.

Model: Y ~ β0 + β1(Sex) + β2(Gestational age in weeks) + β3(Race) + β4(Duplication)

| Coefficient | Estimate | Std.error | t value | p value |
|---|---|---|---|---|
| Sex | 0.46 | 0.23 | 1.99 | 0.050 |
| Gestational age | 0.43 | 0.05 | 7.90 | 8.29E-12 |
| Race (Black) | -0.93 | 1.11 | -0.84 | 0.405 |
| Race (Other) | 0.03 | 1.12 | 0.02 | 0.981 |
| Race (White) | -0.27 | 1.08 | -0.25 | 0.802 |
| Duplication | -0.74 | 0.25 | -2.90 | 0.005 |

Goodness of fit:
Model 1: Y ~ β0 + β1(Sex) + β2(Gestational age in weeks) + β3(Race) + β4(Duplication)
Model 2: Y ~ β0 + β1(Sex) + β2(Gestational age in weeks) + β3(Race)

Model 1 vs. Model 2:

| F | p value |
|---|---|
| 8.39 | 0.005 |

**Table S3: Problems in the first year of life.** Proportion of 3q29Dup participants (n = 31) and typically developing controls (n = 64) reporting problems in the first year of life.

| Condition | 3q29Dup (%, n) | Control (%, n) |
|---|---|---|
| Dehydration | 9.7% (3) | 1.6% (1) |
| Failure to gain weight | 41.9% (13) | 4.7% (3) |
| Feeding problems | 54.8% (17) | 1.6% (1) |
| Hyper-reflexia (overactive reflexes) | 3.2% (1) | 0% (0) |
| Hypo-reflexia (underactive reflexes) | 9.7% (3) | 0% (0) |
| Hypotonia (low muscle tone) | 38.7% (12) | 0% (0) |
| Infection | 19.4% (6) | 1.6% (1) |
| Injury | 3.2% (1) | 1.6% (1) |
| Jaundice | 22.6% (7) | 31.3% (20) |
| Respiratory distress | 29.0% (9) | 0% (0) |
| Unsure | 6.5% (2) | 0% (0) |

**Table S4: Kaplan-Meier time-to-event analysis for developmental milestones.** Time-to-event analysis for developmental milestones across the social-emotional, communication, gross motor, and fine motor domains for 3q29Dup participants and typically developing controls.

| **Developmental Milestone** | **3q29Dup** | | | | | **Control** | | | | |
|---|---|---|---|---|---|---|---|---|---|---|
| | N observations | Mean time to milestone | Median time to milestone | Range | | N observations | Mean time to milestone | Median time to milestone | Range | |
| | | | | Lower | Upper | | | | Lower | Upper |
| **Social-Emotional** | | | | | | | | | | |
| Smile at others | 24 | 6.69 | 5.50 | 1.50 | 33.50 | 63 | 2.90 | 1.50 | 1.50 | 33.50 |
| Play peek-a-boo | 22 | 14.78 | 9.50 | 5.50 | 33.50 | 51 | 6.72 | 5.50 | 1.50 | 15.00 |
| Initiate social interaction by smiling, moving arms, and/or vocalizing | 21 | 9.45 | 5.50 | 1.50 | 33.50 | 56 | 3.91 | 1.50 | 1.50 | 15.00 |
| Cling to caregivers/familiar adults in presence of stranger | 19 | 19.15 | 9.50 | 1.50 | 33.50 | 60 | 9.56 | 9.50 | 1.50 | 42.00 |
| Smile when praised, repeats action for more praise | 20 | 26.17 | 15.00 | 5.50 | 78.00 | 52 | 10.35 | 9.50 | 1.50 | 33.50 |
| Respond affectionately to caregivers (gives hug/kiss when asked) | 21 | 22.47 | 15.00 | 5.50 | 90.00 | 59 | 12.45 | 9.50 | 1.50 | 42.00 |
| Social smile | 17 | 29.92 | 15.00 | 1.50 | 102.00 | 57 | 7.41 | 5.50 | 1.50 | 15.00 |
| **Communication** | | | | | | | | | | |

| | | | | | | | | | | |
|---|---|---|---|---|---|---|---|---|---|---|
| Single syllable utterances (e.g. ma, da) | 23 | 18.73 | 9.50 | 5.50 | 33.50 | 59 | 7.98 | 5.50 | 1.50 | 15.00 |
| First verbal word (for example: go) | 23 | 25.33 | 18.25 | 5.50 | 54.00 | 59 | 11.93 | 9.50 | 5.50 | 21.50 |
| Verbal two word sentences | 25 | 33.26 | 27.50 | 5.50 | 66.00 | 56 | 16.04 | 15.00 | 9.50 | 27.50 |
| Recognizes written letters/numbers | 25 | 66.29 | 66.00 | 15.00 | 102.00 | 53 | 29.09 | 27.50 | 9.50 | 54.00 |
| Phonics-style reading (sounds out words) | 25 | 98.16 | 90.00 | 33.50 | 148.80 | 57 | 55.28 | 54.00 | 9.50 | 78.00 |
| Read whole words (words as an individual unit) | 25 | 104.64 | 102.00 | 54.00 | 149.00 | 59 | 66.30 | 66.00 | 9.50 | 90.00 |
| Write/type using a keyboard | 24 | 105.66 | 102.00 | 54.00 | 149.00 | 59 | 87.79 | 78.00 | 54.00 | 165.00 |
| **Gross motor** | | | | | | | | | | |
| Hold head up on his/her own | 22 | 8.29 | 5.50 | 1.50 | 54.00 | 59 | 3.33 | 1.50 | 1.50 | 9.50 |
| Roll over back to stomach | 21 | 10.75 | 5.50 | 1.50 | 78.00 | 59 | 4.42 | 5.50 | 1.50 | 9.50 |
| Sit when placed | 22 | 10.43 | 9.50 | 5.50 | 21.50 | 61 | 6.44 | 5.50 | 5.50 | 15.00 |
| Crawl on hands and knees | 21 | 13.64 | 15.00 | 5.50 | 27.50 | 62 | 8.62 | 9.50 | 5.50 | 21.50 |
| Walk unassisted | 26 | 18.80 | 15.00 | 15.00 | 33.50 | 62 | 14.24 | 15.00 | 9.50 | 27.50 |
| Climb stairs standing up without help | 23 | 41.30 | 27.50 | 15.00 | 66.00 | 53 | 19.58 | 21.50 | 5.50 | 33.50 |
| Descend stairs without help | 23 | 47.92 | 33.50 | 15.00 | 66.00 | 55 | 23.01 | 21.50 | 15.00 | 54.00 |
| Jump with both feet | 21 | 55.45 | 54.00 | 15.00 | 78.00 | 49 | 24.41 | 21.50 | 15.00 | 54.00 |
| Pedal tricycle | 22 | 64.25 | 54.00 | 21.50 | 114.00 | 51 | 32.50 | 27.50 | 21.50 | 54.00 |
| **Fine motor** | | | | | | | | | | |

| | | | | | | | | | | |
|---|---|---|---|---|---|---|---|---|---|---|
| Look at, reach, and grasp objects placed at a distance | 20 | 11.50 | 5.50 | 1.50 | 33.50 | 52 | 6.49 | 5.50 | 1.50 | 42.00 |
| Transfer object between hands | 22 | 14.21 | 15.00 | 5.50 | 33.50 | 49 | 7.68 | 5.50 | 1.50 | 21.50 |
| Isolate index finger to point | 18 | 40.43 | 21.50 | 9.50 | 78.00 | 41 | 10.22 | 9.50 | 5.50 | 15.00 |
| Clap hands | 22 | 18.38 | 15.00 | 5.50 | 42.00 | 56 | 9.04 | 9.50 | 5.50 | 15.00 |
| Deliberately release object to a container | 17 | 24.76 | 21.50 | 5.50 | 90.00 | 45 | 10.79 | 9.50 | 5.50 | 33.50 |
| Hit two objects together | 19 | 19.94 | 15.00 | 5.50 | 78.00 | 51 | 9.85 | 9.50 | 5.50 | 21.50 |
| Acquired pincer grasp | 14 | 35.09 | 21.50 | 5.50 | 90.00 | 45 | 11.34 | 9.50 | 5.50 | 21.50 |
| Hold and drink from open cup unassisted | 23 | 40.64 | 21.50 | 15.00 | 90.00 | 53 | 17.35 | 15.00 | 5.50 | 42.00 |
| Turn knobs | 21 | 41.60 | 42.00 | 15.00 | 102.00 | 46 | 21.64 | 21.50 | 9.50 | 33.50 |
| Stack and balance blocks | 23 | 36.98 | 27.50 | 15.00 | 90.00 | 49 | 18.15 | 15.00 | 9.50 | 42.00 |

**Table S5: Learning problems.** Proportion of 3q29Dup participants (n = 31) and typically developing controls (n = 64) reporting problems in the first year of life.

| Condition | 3q29Dup (%, n) | Control (%, n) |
|---|---|---|
| Auditory processing disorder | 3.2% (1) | 0% (0) |
| Autism | 38.7% (12) | 0% (0) |
| Dyscalculia | 0% (0) | 0% (0) |
| Dyslexia | 3.2% (1) | 0% (0) |
| Dysphasia/Aphasia | 3.2% (1) | 0% (0) |
| Global developmental delay | 41.9% (13) | 0% (0) |
| Language (receptive) delay (problems understanding language) | 29.0% (9) | 0% (0) |
| Learning disability in math | 25.8% (8) | 0% (0) |
| Learning disability in reading | 25.8% (8) | 3.1% (2) |
| Intellectual disability (mild, moderate, severe) | 6.5% (2) | 0% (0) |
| Non-verbal learning disability | 3.2% (1) | 0% (0) |
| Short term memory problems | 0% (0) | 0% (0) |
| Speech (expressive) delay (problems getting words out) | 54.8% (17) | 3.1% (2) |
| Verbal apraxia/dyspraxia | 3.2% (1) | 0% (0) |
| Visual processing deficits | 3.2% (1) | 0% (0) |
| Writing disability | 16.1% (5) | 0% (0) |
| Unsure | 0% (0) | 1.6% (1) |

**Table S6: Gastrointestinal problems.** Proportion of 3q29Dup participants (n = 31) and typically developing controls (n = 64) reporting gastrointestinal problems.

| Condition | 3q29Dup (%, n) | Control (%, n) |
|---|---|---|
| Anterior displaced anus | 0% (0) | 0% (0) |
| Autistic enterocolitis | 0% (0) | 0% (0) |
| Barrett's esophagus | 0% (0) | 0% (0) |
| Chronic constipation | 35.5% (11) | 7.8% (5) |
| Chronic diarrhea | 6.5% (2) | 4.7% (3) |
| Diaphragmatic hernia | 0% (0) | 0% (0) |
| Dysphagia (difficulty swallowing) | 16.1% (5) | 0% (0) |
| Feeding problems | 38.7% (12) | 1.6% (1) |
| Gastroesophageal reflux (GERD) | 12.9% (4) | 6.3% (4) |
| Hiatial hernia | 0% (0) | 0% (0) |
| Inflammatory bowel disease (Chron's disease, Ulcerative colitis) | 3.2% (1) | 0% (0) |
| Intestinal malrotation | 0% (0) | 0% (0) |
| Irritable bowel syndrome (IBS) | 0% (0) | 0% (0) |
| Peptic ulcers | 0% (0) | 0% (0) |
| Pyloric stenosis | 0% (0) | 0% (0) |
| Silent Reflux | 9.7% (3) | 3.1% (2) |
| Unsure | 16.1% (5) | 0% (0) |

**Table S7: Psychiatric diagnoses.** Proportion of 3q29Dup participants (n = 31) and typically developing controls (n = 64) reporting psychiatric diagnoses.

| Condition | 3q29Dup (%, n) | Control (%, n) |
|---|---|---|
| Addiction | 0% (0) | 0% (0) |
| Anxiety disorder | 32.3% (10) | 6.3% (4) |
| Bipolar/manic depression | 0% (0) | 0% (0) |
| Conduct disorder | 6.5% (2) | 0% (0) |
| Depression | 16.1% (5) | 6.3% (4) |
| Oppositional defiant disorder | 6.5% (2) | 0% (0) |
| Panic attacks | 6.5% (2) | 3.1% (2) |
| Schizophrenia | 0% (0) | 0% (0) |
| Unsure | 12.9% (4) | 0% (0) |

**Table S8: SRS score comparison stratified by ASD status and sex.** Comparison of mean scores on the SRS between study participants with 3q29Del stratified by ASD status and sex to controls (mean ± SD = 45.91 ± 7.97). P values were calculated using simple linear regression, adjusting for age and race.

| | Mean ± SD | P value |
|---|---|---|
| Sex | | |
| Male control | 45.80 ± 6.38 | - |
| Male 3q29Dup | 82.00 ± 22.91 | 1.83E-07 |
| Female control | 46.04 ± 9.62 | - |
| Female 3q29Dup | 77.11 ± 17.51 | 2.22E-07 |
| ASD status | | |
| Control | 45.91 ± 7.97 | - |
| No ASD diagnosis 3q29Dup | 73.67 ± 20.60 | 2.93E-09 |
| ASD diagnosis 3q29Dup | 87.17 ± 14.91 | 3.29E-12 |

**Table S9: Heart defects.** Proportion of 3q29Dup participants (n = 31) and typically developing controls (n = 64) reporting congenital heart defects.

| Heart defect | 3q29Dup (%, n) | Control (%, n) |
|---|---|---|
| Aortic valvar stenosis | 0% (0) | 0% (0) |
| Aortic valve regurgitation | 0% (0) | 0% (0) |
| Atrial septal defect | 3.2% (1) | 0% (0) |
| Atrioventricular septal defect (or atrioventricular canal defect) | 0% (0) | 0% (0) |
| Coarctation of the aorta | 0% (0) | 0% (0) |
| Ebstein's anomaly | 0% (0) | 0% (0) |
| Hypoplastic left heart syndrome | 0% (0) | 0% (0) |
| Interrupted aortic arch/ventricular septic defect | 0% (0) | 0% (0) |
| Mitral valve regurgitation | 0% (0) | 0% (0) |
| Patent ductus arteriosus | 0% (0) | 0% (0) |
| Pulmonary atresia | 0% (0) | 0% (0) |
| Pulmonary valvar stenosis | 0% (0) | 0% (0) |
| Pulmonary valve regurgitation | 0% (0) | 0% (0) |
| Single ventricle anomalies | 0% (0) | 0% (0) |
| Tetralogy of Fallot | 0% (0) | 0% (0) |
| Total anomalous pulmonary venous return | 0% (0) | 0% (0) |
| Transposition of the great arteries | 0% (0) | 0% (0) |
| Tricuspid atresia | 3.2% (1) | 0% (0) |
| Tricuspid valve regurgitation | 0% (0) | 0% (0) |
| Truncus arteriosus | 0% (0) | 0% (0) |
| Vascular rings | 0% (0) | 0% (0) |
| Ventricular septal defect | 0% (0) | 0% (0) |
| Unsure | 12.9% (4) | 0% (0) |

**Table S10: Genitourinary problems.** Proportion of 3q29Dup participants (n = 31) and typically developing controls (n = 64) reporting genitourinary problems.

| Condition | 3q29Dup (%, n) | Control (%, n) |
|---|---|---|
| Ambiguous genitalia | 0% (0) | 0% (0) |
| Bladder diverticulum | 3.2% (1) | 0% (0) |
| Bladder extrophy | 3.2% (1) | 0% (0) |
| Difficulty emptying baldder | 3.2% (1) | 0% (0) |
| Hypospadia | 0% (0) | 0% (0) |
| Large bladder | 0% (0) | 0% (0) |
| Micropenis | 3.2% (1) | 0% (0) |
| Posterior urethral valves | 0% (0) | 0% (0) |
| Small bladder | 0% (0) | 0% (0) |
| Undescended testicle (cryptorchidism) | 0% (0) | 1.6% (1) |
| Ureter reimplantation | 0% (0) | 0% (0) |
| Unsure | 19.4% (6) | 0% (0) |

**Table S11: Renal problems.** Proportion of 3q29Dup participants (n = 31) and typically developing controls (n = 64) reporting renal problems.

| Condition | 3q29Dup (%, n) | Control (%, n) |
|---|---|---|
| Cystic kidney (polycystic kidney) | 0% (0) | 0% (0) |
| Dilated renal pelvis | 0% (0) | 0% (0) |
| Duplicate or extra kidney | 0% (0) | 0% (0) |
| Horseshoe kidney | 0% (0) | 0% (0) |
| Hydronephrosis | 0% (0) | 0% (0) |
| Increased kidney size | 0% (0) | 0% (0) |
| Kidney reflux (vesicoureteral reflux) | 3.2% (1) | 0% (0) |
| Kidney stones | 0% (0) | 0% (0) |
| Malformed kidney | 0% (0) | 1.6% (1) |
| Missing kidney(s) (renal agenesis) | 0% (0) | 0% (0) |
| Recurrent UTIs | 16.1% (5) | 0% (0) |
| Unsure | 9.7% (3) | 0% (0) |

**Table S12: Ear problems.** Proportion of 3q29Dup participants (n = 31) and typically developing controls (n = 64) reporting ear problems.

| Condition | 3q29Dup (%, n) | Control (%, n) |
|---|---|---|
| Dizziness and/or vertigo | 9.7% (3) | 0% (0) |
| Ear pain | 16.1% (5) | 7.8% (5) |
| Meniere's disease | 0% (0) | 0% (0) |
| Recurrent ear infections | 35.5% (11) | 20.3% (13) |
| Tinnitus (ringing in the ear) | 3.2% (1) | 0% (0) |
| Unsure | 9.7% (3) | 0% (0) |

**Table S13: Dental problems.** Proportion of 3q29Dup participants (n = 31) and typically developing controls (n = 64) reporting dental problems.

| Condition | 3q29Dup (%, n) | Control (%, n) |
|---|---|---|
| Cone-shaped teeth | 9.7% (3) | 0% (0) |
| Crowded teeth | 6.5% (2) | 3.1% (2) |
| Enamel hypoplasia | 6.5% (2) | 1.6% (1) |
| Extra teeth | 0% (0) | 1.6% (1) |
| High number of cavities | 25.8% (8) | 6.3% (4) |
| Large gap between two front teeth on top | 22.6% (7) | 6.3% (4) |
| Large teeth | 3.2% (1) | 0% (0) |
| Malocclusion | 3.2% (1) | 1.6% (1) |
| Missing teeth | 3.2% (1) | 1.6% (1) |
| Small teeth | 16.1% (5) | 3.1% (2) |
| Tooth in palate (roof of mouth) | 0% (0) | 1.6% (1) |
| Tooth/teeth extraction (pulled) | 19.4% (6) | 6.3% (4) |
| Weak/soft tooth enamel | 12.9% (4) | 1.6% (1) |
| Widely speced teeth | 6.5% (2) | 1.6% (1) |
| Unsure - dental | 16.1% (5) | 0% (0) |

**Table S14: Clefting.** Proportion of 3q29Dup participants (n = 31) and typically developing controls (n = 64) reporting clefting.

| Condition | 3q29Dup (%, n) | Control (%, n) |
|---|---|---|
| Cleft lip | 0% (0) | 1.6% (1) |
| Cleft palate | 3.2% (1) | 0% (0) |
| Unsure | 6.5% (2) | 0% (0) |

**Table S15: Psychiatric diagnosis comparison between 3q29Dup and 3q29Del.** Comparison of reported psychiatric diagnoses between 3q29Dup participants (n = 31) and 3q29Del participants (n = 117). P values were calculated using chi-square test or Fisher's exact test.

| Diagnosis | 3q29Dup (%, n) | 3q29Del (%, n) | P value |
|---|---|---|---|
| ASD | 38.7% (12) | 29.1% (34) | 0.416 |
| Addiction | 0% (0) | 2.6% (3) | 1.00 |
| Anxiety disorder | 32.3% (10) | 28.2% (33) | 0.048 |
| Bipolar/manic depression | 0% (0) | 5.1% (6) | 0.344 |
| Conduct disorder | 6.5% (2) | 3.4% (4) | 0.606 |
| Depression | 16.1% (5) | 6.0% (7) | 0.130 |
| Oppositional defiant disorder | 6.5% (2) | 4.3% (5) | 0.637 |
| Panic attacks | 6.5% (2) | 12.0% (14) | 0.525 |
| Schizophrenia | 0% (0) | 3.4% (4) | 0.580 |
